# Epidemiology of Cancers in Zambia: A Significant Variation in Cancer Incidence and Prevalence across the Nation

**DOI:** 10.1101/402628

**Authors:** Maybin Kalubula, Heqing Shen, Mpundu Makasa, Longjian Liu

## Abstract

**Background:** Cancers are one of the leading causes of death worldwide. More than two thirds of deaths due to cancers occur in low- and middle-income countries whereZambia belongs. This study therefore sought to assess the epidemiology of cancers in Zambia.

**Methods:** We conducted a retrospective observational study nested on Zambia National Cancer Registry (ZNCR) histopathological and clinical data from 2007 to 2014. Zambia Central Statistics Office (CSO)demographic datawere used to calculate prevalence and incidence rates of cancers. Age-adjusted rates and case fatality rates were estimated using standard methods. We used a Poisson Approximation for calculating 95% confidence intervals (CI).

**Results:** The top seven most cancer prevalent districts in Zambia have been Luangwa, Kabwe, Lusaka, Monze, Mongu, Katete and Chipata. Cervical cancer, prostate cancer, breast cancer and Kaposi’s sarcoma were the top four most prevalent cancers as well as major causes of cancer related deaths in Zambia. Standardised Incidence Rates and 95% CI for the top four cancers were: cervix uteri (186.3; CI = 181.77 – 190.83), prostate (60.03; CI = 57.03 – 63.03), breast (38.08; CI = 36.0 – 40.16) and Kaposi’s sarcoma (26.18; CI = 25.14 – 27.22).CFR were: Leukaemia (38.1%); pancreatic cancer (36.3%); lung cancer (33.3%); and brain, nervous system (30.2%). Cancers were associated with HIV with *p*-value of 0.000 and Pearson correlation coefficient of 0.818.

**Conclusions:** The widespread distribution of cancers with high prevalence in the southern zone has been perpetrated by lifestyle and sexual culture as well as geography. Intensifying cancer screening and early detection countrywide as well as changing the lifestyle and sexual culture would greatly help in the reduction of cancer cases in Zambia.

## INTRODUCTION

The global burden of cancers has been on the increase over the past few decades despite some remarkable advances in early cancer detection, treatment and prevention [1]. Cancers have continued to claim human lives and are one of the leading causes of death worldwide hence the realization by many organizations and researchers to make advancements inconventional medicine as well as anticancer nanomedicine and nanotechnology to broaden the spectrum of combating the scourge [2-4]. In the year 2013, cancers caused over 8 million deaths worldwide and have moved from the third leading cause of death in 1990 to the second leading cause behind cardiovascular diseases in 2013[5-7]. More than two thirds of deaths due to cancers occur in low- and middle-income countries [8]. Low-income countries reported approximately 51% of all cancers globally in the year 1975 but this proportion steadily increased to 55% in 2007. It is estimated that by 2050, low-incomecountries will account for 61% of all cancers globally [9-11].

The global increase in the number of cancer cases is due to multiple factors such as lifestyle trends that are associated with economic development and the increasing adoption of risk behaviour such as consumption of unhealthy diets, lack of physical exercise, harmful use of alcohol and tobacco as well as exposures to environmental pollutants [12-15]. Thirdworld countries have 26% cancers attributable to infection while developed countries have 8% which is one third of cancers attributable to infection in thirdworld countries [16]. The oncogenic infections that have been linked to these cancers are Human Papilloma Virus (HPV) [17], Hepatitis B Virus (HBV) [18], *Helicobacter pylori* (*H. pylori*) [19], Human Herpes Virus (HHV8) [20], and Epstein Barr Virus (EBV) [21].

Although cancer incidence has been increasing in every part of the world, there are huge inequalities between developed and third world countries. Incidence rates remain highest in developed countries, but mortality is relatively much higher in third world countries due to lack of early detection and access to treatment facilities[8]. Infections due to human papillomavirus and hepatitis B and C viruses significantly contribute to the burden of cervical and liver cancers respectively on the African Continent [8]. The most common cancers in the African region are cervical, breast, liver and prostate as well as Kaposi’s sarcoma and non-Hodgkin’s lymphoma [8].

Zambia is one of the sub-Saharan African countries that have not been spared by the increasing burden of cancers. Many lives can be saved if appropriate investment is made in raising public awareness on the early signs and symptoms of common cancers as well as implementation of early detection strategies [8]. This study therefore sought to assess the epidemiology of cancers in Zambia with a view of finding amicable solution.

## MATERIALS AND METHODS

### Study site

The Republic of Zambia is located in the southern zone of the African continent and lies between latitudes 8° and 18° south, and longitudes 22° and 34° east. The country has a total geographical area of 753,612km^2^ [22].

### Study design

We conducted a retrospective observational study nested on Zambia National Cancer Registry (ZNCR) histopathological and clinical data from 2007 to 2014. ZNCR collects and keeps medical records of all diagnosed cancers in Zambia. The collected information include core variables such as patients personal details (names, age, sex, date of birth and residential address at diagnosis), hospital details (hospital, consultant patient unit number), diagnostics, tumour and treatment details (site of primary, morphology, laterality, stage, grade of tumour, basis of diagnosis, date of diagnosis, treatment indicators) and death details (alive or dead, date of death, cause(s) and place of death).

All health facilities countrywide submit reports of all diagnosed cancers to ZNCR. Medical records collected in form of hard copies are entered and stored in the ZNCR main database by data entry clerks. ZNCR ensures that cancer registrations data is stored securely, backed up and only accessible to authorised cancer registry staff. All cancer notification forms or books are filed and locked in a secure place. Information request for data on cancers are made in writing to the Registrar and the Permanent Secretary at the Ministry of Health for approval. This study was therefore approved by both the University of Zambia Biomedical Research Ethics Committee (UNZABREC) and the National Health Research Authority of Zambia.

Zambia Central Statistics Office (CSO) provided us with shapes files for provinces and districts as well as demographic data which we used to determine prevalence and Age-specific rates while World Standard Population [23] was used to determine Standardised Incidence Rates (SIR). 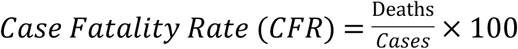 We used a Poisson Approximation for calculating 95% confidence intervals. The 95% confidence interval for the age standardised rate is given by:

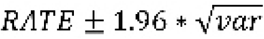

Where the variance for the age standardised rate is given by:

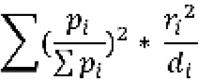

*d_i_* is the number of events in age group *i* in the study population

*r_i_* is the incidence rate in the study population for the persons in age group *i*

*𝒫i* is the number of persons in age group *i* in the standard population.

Mapping of the distribution of cancers at district and provincial levels were done using Geographical Information System (ArcGIS) version 10.3.1, Redlands, CA. Districts such as Ikelenge, Zimba and Mafinga were still part of the original districts namely, Mwinilunga, Kalomo and Isoka respectively during the study period hence we considered cancer rates of original districts in these new districts.

### Sampling

Since our study was basically registry based, we considered all cancer cases in the Zambia National Cancer Registry database from 2007 to 2014. We however excluded cancer cases not belonging to any of the ten provinces in Zambia (i.e. not coded cases from any region in Zambia) for the purpose of GIS mapping which required that cancer cases be linked to districts and provinces of origin. In addition, we sampled and surveyed ten districts across Zambia to assess the cancer situation and determine some risk factors contributing to escalating cancer cases.

### Statistical Analyses

We used SPSS version 21 in our data analyses. A linear regression model was used to determine the association between cancers and HIV while Pearson Correlation was used to determine the correlation coefficient. We used a Poisson Approximation for calculating 95% confidence intervals.

## RESULTS

### Distribution of cancers by region and province

Cancers are widely distributed in Zambiawith high prevalence concentrated in the southern zone comprising Eastern, Central, Lusaka, Western and Southern Provinces. The southern zone can be divided into high prevalence and medium prevalence regions. Eastern and Lusaka Provinces form the high prevalence region while Central, Southern and Western Provinces form the medium prevalence region. The northern zone comprising North Western, Copperbelt, Luapula, Northern and Muchinga Provinces has relatively low prevalence of cancers. **Figure 1** shows the distribution of all types of cancers in Zambia by region and province per 100,000 population.

By sex, cancers have been more prevalent in females than in males in all the ten provinces with mean sex ratio F/M = 1.85. **Table 1** shows the prevalence of cancers by province and sex. Cancers in males were more prevalent in Lusaka, Eastern,Central and North Western Provinces. Cancers in females were more prevalent in Lusaka, Eastern, Central, Southern,Western and Copperbelt Provinces. **Figure 2** shows details of provincial distribution of cancers by sex.

**Table 1:**
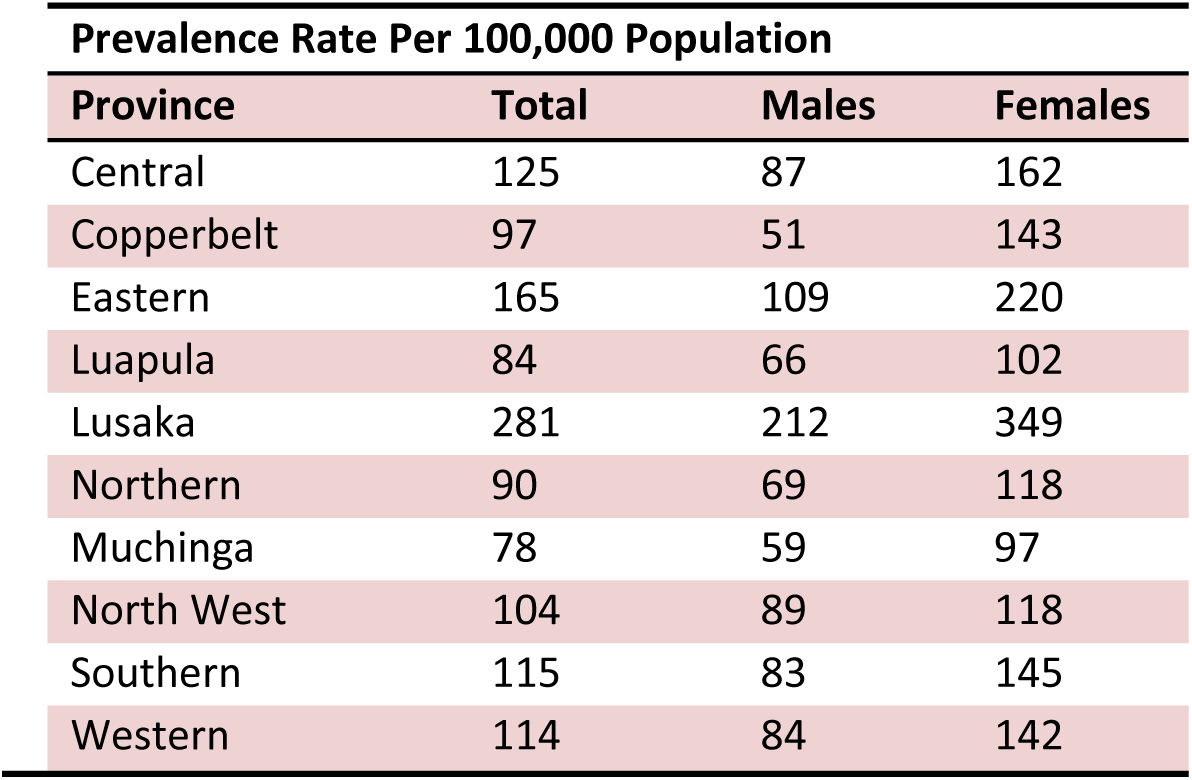
Distribution of cancers (all types) in Zambia by Province

### Distribution of cancers by district

The distribution of cancers at district level is similar to the zonal pattern at provincial level. **Figure 3** shows the geographical distribution of cancers by district in Zambia. Some districts such as Mafinga, Ikelenge and Zimba were still part of the original districts namely Isoka, Mwinilunga and Kalomo respectively at the beginning of our research period hence are shown as combined districts in Table 2. The prevalence of cancers in the original districts reflects cancer prevalence in the new districts.

**Table 2:** Ranked prevalence of cancers (all types) by district per 100,000 population

| District | Total | Males | Females | District | Total | Males | Females |
| --- | --- | --- | --- | --- | --- | --- | --- |
| Luangwa | 1014 | 766 | 1257 | Chinsali | 90 | 52 | 127 |
| Kabwe | 379 | 251 | 499 | Mkushi | 88 | 57 | 120 |
| Lusaka | 311 | 233 | 387 | Mwense | 88 | 76 | 99 |
| Monze | 282 | 157 | 404 | Luwingu | 87 | 360 | 114 |
| Mongu | 260 | 176 | 337 | Kitwe | 82 | 52 | 113 |
| Katete | 229 | 142 | 313 | Kaoma | 80 | 62 | 96 |
| Chipata | 213 | 133 | 291 | Sesheke | 78 | 64 | 91 |
| Kasempa | 193 | 176 | 210 | Isoka/ Mafinga | 77 | 62 | 91 |
| Kasama | 168 | 118 | 217 | Kafue | 77 | 66 | 88 |
| Ndola | 165 | 76 | 252 | Lukulu | 77 | 37 | 114 |
| Mansa | 165 | 110 | 217 | Chadiza | 74 | 49 | 99 |
| Choma | 158 | 126 | 189 | Samfya | 70 | 74 | 67 |
| Luanshya | 156 | 77 | 234 | Siavonga | 70 | 49 | 90 |
| Chongwe | 155 | 123 | 187 | Namwala | 69 | 66 | 71 |
| Petauke | 153 | 102 | 202 | Solwezi | 65 | 60 | 71 |
| Mumbwa | 144 | 116 | 171 | KapiriMposhi | 61 | 32 | 89 |
| Mbala | 142 | 96 | 187 | Chibombo | 56 | 45 | 68 |
| Livingstone | 138 | 118 | 157 | Nakonde | 53 | 36 | 70 |
| Zambezi | 135 | 116 | 155 | Sinazongwe | 48 | 49 | 47 |
| Mazabuka | 126 | 86 | 165 | Chama | 48 | 47 | 48 |
| Lundazi | 120 | 86 | 153 | Kalomo/ Zimba | 45 | 34 | 47 |
| Gwembe | 113 | 80 | 145 | Mpongwe | 44 | 19 | 70 |
| Mambwe | 112 | 110 | 113 | Nchelenge | 43 | 30 | 55 |
| Kalabo | 111 | 91 | 130 | Chililabombwe | 40 | 19 | 61 |
| Mufumbwe | 110 | 97 | 124 | Kalulushi | 35 | 13 | 57 |
| Mwinilunga/Ikelenge | 107 | 73 | 141 | Milenge | 29 | 20 | 38 |
| Nyimba | 107 | 72 | 141 | Mpulungu | 29 | 14 | 43 |
| Kabompo | 106 | 97 | 116 | Mungwi | 27 | 19 | 35 |
| Serenje | 105 | 78 | 132 | Kaputa | 26 | 16 | 35 |
| Chavuma | 105 | 102 | 108 | Shang'ombo | 19 | 18 | 20 |
| Mufulira | 104 | 60 | 148 | Chilubi | 18 | 14 | 22 |
| Mpika | 102 | 82 | 122 | Masaiti | 17 | 17 | 17 |
| Mporokoso | 100 | 77 | 123 | Lufwanyama | 16 | 2 | 31 |
| Kawambwa | 95 | 78 | 110 | ItezHITEZHI | 16 | 17 | 14 |
| Senanga | 93 | 84 | 101 | Kazungula | 14 | 13 | 16 |
| Chingola | 93 | 53 | 133 | Chiengi | 12 | 9 | 15 |

Luangwa has been the most cancer prevalent district in Zambia with the rate of 1014 per 100,000 population, seconded by Kabwe with the prevalence rate of 379 per 100,000 population. Lusaka has been the third district with prevalence rate of 311 per 100,000, Monze ranks fourth with the rate of 282 per 100,000, Mongu ranked fifth with prevalence rate of 260 per 100,000, Katete ranked sixth with prevalence rate of 229 and Chipata district ranked seventh with prevalence of 213 per 100,000 population. All these seven districts are from the southern zone.

The top four cancer prevalent districts in the northern zone are Kasempa with prevalence rate of 193 per 100,000, Kasama with prevalence rate of 168 per 100,000, Ndola with prevalence rate of 165 per 100,000 and Mansa with prevalence rate of 165 per 100,000 population. The least four cancer prevalent districts in Zambia were Lufwanyama with prevalence rate of16per 100,000,Itezhitezhi with prevalence rate of 16 per 100,000, Kazungula with prevalence rate of 14 per 100,000 and Chienge with prevalence rate of 12 per 100,000 population during the study period. **Table 2** shows details of ranked prevalence of cancers by district per 100,000 population.

With a few exceptions of districts such as Luwingu, Sinazongwe and Samfya where cancers are more prevalent in males than in females, most districts in Zambia have high cancer prevalence in females than in males. The trend in the total district pattern (see Figure 3)remains the same as cancer distribution by sex at district level (see Figure 4). The detailed geographical distribution of cancers by sex at district level is shown in **Figure 4**.

## Morbidity and mortality of cancers in Zambia

During our study period from 2007 to 2014, a total of 21,512 cancer cases have been notified to ZNCR of which 7,560 (35.14%) were males and 13,952 (64.86%) were females. The actual notified cases could slightly be higher than these figures but due to missing or incomplete information, some cases could not be linked to districts and provinces of origin as we employed Geographical Information System (ArcGIS) version 10.3.1 to map cancers. Cervical cancer is the most prevalent cancer in Zambia followed by prostate cancer, breast cancer and Kaposi’s sarcoma in that order. The prevalence rate of cervical cancer has been 97.1 per 100,000 females and represents 34.3% of all cancers in Zambia. In proportional terms, Kaposi’s sarcoma ranks second and represents 13.3%, prostate cancer ranks third and represents 7.7% while breast cancer ranks fourth and represents 6.8%. Myeloma and Other Pharynx are the least prevalent among other cancers, each representing 0.2% of all cancers in Zambia. **Table 3** shows details of prevalence and proportions of cancers in Zambia from 2007 to 2014.

**Table 3:** The Prevalence and proportions of cancers in Zambia, 2007 - 2014

| Cancer Type | Cases Both Sexes | Prevalence per 100,000 pop. | Male |  | Female |  | Both Sexes Proportion (%) |
| --- | --- | --- | --- | --- | --- | --- | --- |
|  |  |  | Cases | Proportion (%) | Cases | Proportion (%) |  |
| Cervix Uteri | 7389 | 97.1 |  |  | 7389 | 53 | 34.3 |
| Kaposi's Sarcoma | 2891 | 19.2 | 1759 | 23.3 | 1132 | 8.1 | 13.4 |
| Prostate | 1651 | 22.1 | 1651 | 21.8 |  |  | 7.7 |
| Breast | 1469 | 19.3 | 41 | 0.5 | 1428 | 10.2 | 6.8 |
| Eye | 909 | 6.0 | 431 | 5.7 | 478 | 3.4 | 4.2 |
| Non-Hodgkin Lymphoma | 665 | 4.4 | 358 | 4.7 | 307 | 2.2 | 3.1 |
| Oesophagus | 581 | 3.9 | 365 | 4.8 | 216 | 1.5 | 2.7 |
| Large Bowel | 539 | 3.6 | 278 | 3.7 | 261 | 1.9 | 2.5 |
| Bladder | 478 | 3.2 | 261 | 3.5 | 217 | 1.6 | 2.2 |
| Liver | 460 | 3.1 | 283 | 3.7 | 177 | 1.3 | 2.1 |
| Other Skin | 406 | 2.7 | 192 | 2.5 | 214 | 1.5 | 1.9 |
| Stomach | 395 | 2.6 | 210 | 2.8 | 185 | 1.3 | 1.8 |
| Bone | 252 | 1.7 | 130 | 1.7 | 122 | 0.9 | 1.2 |
| Vulva/Vagina | 246 | 3.2 |  |  | 246 | 1.8 | 1.1 |
| Ovary | 238 | 3.1 |  |  | 238 | 1.7 | 1.1 |
| Kidney | 171 | 1.1 | 80 | 1.1 | 91 | 0.7 | 0.8 |
| Oral Cavity | 168 | 1.1 | 95 | 1.3 | 73 | 0.5 | 0.8 |
| Penis | 166 | 2.2 | 166 | 2.2 |  |  | 0.8 |
| Hodgkin Disease | 151 | 1.0 | 99 | 1.3 | 52 | 0.4 | 0.7 |
| Melanoma of the skin | 145 | 1.0 | 56 | 0.7 | 89 | 0.6 | 0.7 |
| Lung | 135 | 0.9 | 90 | 1.2 | 45 | 0.3 | 0.6 |
| Brain, Nervous System | 126 | 0.8 | 69 | 0.9 | 57 | 0.4 | 0.6 |
| Leukaemia | 126 | 0.8 | 79 | 1 | 47 | 0.3 | 0.6 |
| Larynx | 125 | 0.8 | 102 | 1.3 | 23 | 0.2 | 0.6 |
| Uterus | 115 | 1.5 |  |  | 115 | 0.8 | 0.5 |
| Pancreas | 102 | 0.7 | 61 | 0.8 | 41 | 0.3 | 0.5 |
| Corpus Uteri | 95 | 0.6 |  |  | 95 | 0.7 | 0.4 |
| Thyroid | 92 | 0.6 | 30 | 0.4 | 62 | 0.4 | 0.4 |
| Nasopharynx | 85 | 0.6 | 51 | 0.7 | 34 | 0.2 | 0.4 |
| Myeloma | 45 | 0.3 | 27 | 0.4 | 18 | 0.1 | 0.2 |
| Other Pharynx | 42 | 0.3 | 29 | 0.4 | 13 | 0.1 | 0.2 |
| Others | 1054 | 7.0 | 567 | 7.5 | 487 | 3.5 | 4.9 |
| **Totals** | **21512** | **142.8** | **7560** | **100** | **13952** | **100** | **100** |

Age specific rates, Standardised Incidence Rates (SIR) and the 95% confidence intervals (CI) for all cancers in Zambia are shown in **Table 4**. Rates were adjusted using the world standard population [23]. The standardised incidence ratesand 95% confidence intervalsfor the top four cancers in Zambia were: cervix uteri (186.3; 95% CI = 181.77 – 190.83), prostate (60.03; 95% CI = 57.03 – 63.03), breast (38.08; 95% CI = 36.0 – 40.16) and Kaposi’s sarcoma (26.18;95% CI = 25.14 – 27.22). Peaks of age specific rates were in the age range 40 – 49 years for cervix uteri, 60 – 69 years for prostate cancer, and 40 – 49 years for breast cancer and 30 – 39 years for Kaposi’s sarcoma.

**Table 4:** Standardised Incidence Rates (SIR) for all cancers in Zambia, 2007 - 2014

| CANCER TYPE | Age-specific Incidence Rate per 100,000 population |  |  |  |  |  |  |  |  | SIR all ages | 95% CI |  |
| --- | --- | --- | --- | --- | --- | --- | --- | --- | --- | --- | --- | --- |
|  | 0-9yr | 10-19 | 20-29 | 30-39 | 40-49 | 50-59 | 60-69 | 70-79 | 80+ |  | Lower | Upper |
| Cervix Uteri | 0 | 0.07 | 5.5 | 23.41 | 49.68 | 46.11 | 39.78 | 14.09 | 7.67 | **186.3** | 181.77 | 190.83 |
| Prostate | 0 | 0 | 0 | 0.07 | 0.89 | 5.52 | 22.83 | 21.92 | 8.8 | **60.03** | 57.03 | 63.03 |
| Breast | 0 | 0.03 | 1.29 | 4.03 | 9.46 | 8.53 | 9.09 | 4.38 | 1.26 | **38.08** | 36 | 40.16 |
| Kaposi's Sarcoma | 0.35 | 0.65 | 3.58 | 7.66 | 7.42 | 3.17 | 1.82 | 0.64 | 0.88 | **26.18** | 25.14 | 27.22 |
| Oesophagus | 0.01 | 0.01 | 0.11 | 0.64 | 1.6 | 1.67 | 2.49 | 1.45 | 0.4 | **8.38** | 7.66 | 8.38 |
| Eye | 0.7 | 0.1 | 0.57 | 1.97 | 2.27 | 1.06 | 0.85 | 0.25 | 0.52 | **8.29** | 7.69 | 8.89 |
| Bladder | 0.01 | 0.02 | 0.12 | 0.34 | 0.78 | 1.36 | 2.94 | 1.41 | 0.46 | **7.45** | 6.74 | 8.16 |
| Large Bowel | 0 | 0.03 | 0.32 | 0.64 | 1.63 | 1.53 | 1.93 | 0.87 | 0.24 | **7.19** | 6.54 | 7.84 |
| Non-Hodgkin Lymphoma | 0.31 | 0.45 | 0.51 | 0.8 | 1.46 | 1.33 | 1.28 | 0.48 | 0.2 | **6.82** | 6.23 | 7.41 |
| Vulva/Vagina | 0 | 0.01 | 0.31 | 0.8 | 1.61 | 1.22 | 1.23 | 0.46 | 0.3 | **5.94** | 5.14 | 6.74 |
| Liver | 0.02 | 0.09 | 0.27 | 0.56 | 1.21 | 1.22 | 1.3 | 0.83 | 0.36 | **5.86** | 5.28 | 6.44 |
| Ovary | 0.01 | 0.14 | 0.41 | 0.47 | 1.56 | 1.34 | 0.98 | 0.62 | 0.14 | **5.66** | 4.88 | 6.44 |
| Stomach | 0.01 | 0.02 | 0.12 | 0.41 | 0.93 | 1.23 | 1.53 | 1.04 | 0.32 | **5.61** | 5.02 | 6.2 |
| Other Skin | 0.05 | 0.08 | 0.28 | 0.53 | 0.81 | 1 | 1.26 | 0.69 | 0.38 | **5.07** | 4.53 | 5.61 |
| Penis | 0.02 | 0 | 0.03 | 0.26 | 0.98 | 1.28 | 1.38 | 0.58 | 0.41 | **4.93** | 4.13 | 5.73 |
| Uterus | 0 | 0.01 | 0.12 | 0.26 | 1.04 | 0.61 | 0.64 | 0.1 | 0.11 | **2.89** | 2.33 | 3.45 |
| Corpus Uteri | 0 | 0 | 0.06 | 0.12 | 0.42 | 0.85 | 1.15 | 0.2 | 0.07 | **2.87** | 2.26 | 3.48 |
| Melanoma of the skin | 0 | 0.02 | 0.02 | 0.1 | 0.28 | 0.48 | 0.88 | 0.39 | 0.09 | **2.26** | 1.87 | 2.65 |
| Bone | 0.1 | 0.38 | 0.25 | 0.22 | 0.36 | 0.22 | 0.52 | 0.12 | 0.07 | **2.25** | 1.92 | 2.58 |
| Oral Cavity | 0.04 | 0.05 | 0.08 | 0.13 | 0.36 | 0.55 | 0.63 | 0.34 | 0.07 | **2.24** | 1.87 | 2.61 |
| Larynx | 0.01 | 0 | 0.01 | 0.01 | 0.24 | 0.45 | 0.99 | 0.35 | 0.07 | **2.14** | 1.75 | 2.53 |
| Lung | 0 | 0 | 0.03 | 0.07 | 0.2 | 0.55 | 0.72 | 0.39 | 0.16 | **2.12** | 1.75 | 2.49 |
| Pancreas | 0 | 0 | 0.02 | 0.06 | 0.34 | 0.5 | 0.27 | 0.19 | 0.06 | **1.45** | 1.16 | 1.75 |
| Hodgkin Disease | 0.08 | 0.19 | 0.13 | 0.15 | 0.32 | 0.19 | 0.2 | 0.05 | 0.04 | **1.34** | 1.09 | 1.59 |
| Thyroid | 0 | 0.04 | 0.07 | 0.06 | 0.2 | 0.27 | 0.38 | 0.11 | 0.09 | **1.22** | 0.95 | 1.49 |
| Kidney | 0.45 | 0.09 | 0.01 | 0.1 | 0.13 | 0.12 | 0.11 | 0.07 | 0.07 | **1.16** | 0.95 | 1.37 |
| Brain, Nervous System | 0.16 | 0.08 | 0.1 | 0.08 | 0.23 | 0.2 | 0.18 | 0.09 | 0.02 | **1.13** | 0.9 | 1.36 |
| Nasopharynx | 0.03 | 0.05 | 0.04 | 0.11 | 0.21 | 0.25 | 0.29 | 0.02 | 0 | **0.99** | 0.75 | 1.23 |
| Leukaemia | 0.24 | 0.21 | 0.04 | 0.04 | 0.09 | 0.06 | 0.04 | 0.05 | 0 | **0.78** | 0.62 | 0.94 |
| Other Pharynx | 0 | 0.01 | 0.01 | 0.01 | 0.07 | 0.11 | 0.29 | 0.09 | 0.06 | **0.65** | 0.58 | 0.72 |
| Myeloma | 0 | 0.01 | 0.01 | 0.04 | 0.07 | 0.28 | 0.16 | 0.04 | 0.02 | **0.63** | 0.43 | 0.83 |
| Others | 0.35 | 0.52 | 0.84 | 1.07 | 1.98 | 2.47 | 2.88 | 1.32 | 0.68 | **12.11** | 11.29 | 12.93 |

Like other diseases, cancers have been classified in the International Classification of Diseases known as ICD10. Table 5 shows details of ICD10 codes for cancers, their mortality and case fatality rates (CFR). Leukaemia (38.1%) and pancreatic cancer (36.3%)have the highest CFR among cancers in Zambia followed by lung cancer (33.3%) and brain, nervous system (30.2%).

**Table 5:** Cancer classification and their mortality and CFR in Zambia, 2007 - 2014

| **Cancer Type** | **International Classification of Diseases (ICD10)** | **Cases** | **Deaths** | **Case Fatality Rate (%)** |
| --- | --- | --- | --- | --- |
| Leukaemia | C91-C95 | 126 | 48 | 38.1 |
| Pancreas | C25 | 102 | 37 | 36.3 |
| Lung | C33-C34 | 135 | 45 | 33.3 |
| Brain, Nervous System | C70-C72 | 126 | 38 | 30.2 |
| Kidney | C64 | 171 | 38 | 22.2 |
| Liver | C22 | 460 | 91 | 19.8 |
| Stomach | C16 | 395 | 78 | 19.7 |
| Oesophagus | C15 | 581 | 113 | 19.4 |
| Bladder | C67 | 478 | 90 | 18.8 |
| Ovary | C56 | 238 | 44 | 18.5 |
| Myeloma | C90 | 45 | 8 | 17.8 |
| Nasopharynx | C11 | 85 | 15 | 17.6 |
| Corpus Uteri | C54 | 95 | 16 | 16.8 |
| Bone | C40-C41 | 252 | 40 | 15.9 |
| Melanoma of the Skin | C43 | 145 | 21 | 14.5 |
| Non-Hodgkin Lymphoma | C82,C85,C96 | 665 | 89 | 13.4 |
| Prostate | C61 | 1651 | 215 | 13.0 |
| Large Bowel | C18-C21 | 539 | 68 | 12.6 |
| Kaposi's Sarcoma | C46 | 2891 | 332 | 11.5 |
| Thyroid | C73 | 92 | 10 | 10.9 |
| Larynx | C32 | 125 | 12 | 9.6 |
| Hodgkin Disease | C81 | 151 | 14 | 9.3 |
| Uterus | C55 | 115 | 9 | 7.8 |
| Breast | C50 | 1469 | 113 | 7.7 |
| Cervix Uteri | C53 | 7389 | 543 | 7.3 |
| Penis | C60 | 166 | 12 | 7.2 |
| Oral Cavity | C00-C06 | 169 | 12 | 7.1 |
| Other Skin | C44 | 406 | 25 | 6.2 |
| Eye | C69 | 909 | 53 | 5.8 |
| Vulva/Vagina | C51-C52 | 246 | 12 | 4.9 |
| Other Pharynx | C09-C10, C12-C14 | 42 | 2 | 4.8 |
| Others | C07-C08, C17,... | 1054 | 154 | 14.6 |
| **Totals** |  | **21512** | **2397** | **11.1** |

Although CFR has been very high in Leukaemia,pancreatic cancer,lung cancer and cancers of the brain and nervous system, cancer mortality rates have been high in cervical cancer, prostate cancer, Kaposis’s sarcoma, breast cancer and oesophageal cancer in that order. During our study period 2007 to 2014, there were 543 cervical cancer deaths. Prostate cancer ranked second with 215 deaths followed by Kaposis’ sarcoma with 332 deaths, then breast cancer and oesophagus cancer each with 113 deaths. Note: mortality rate for prostate cancer is higher than that of Kaposi’s sarcoma despite the fact that Kaposi’s sarcoma reported higher absolute number of 332 deaths while prostate cancer reported 215 deaths. The same principle applies for breast cancer and oesophagus cancer where breast cancer has a higher mortality rate than oesophageal cancer despite having the same absolute death figures. The reason is that the target population (denominator) for Kaposi’ssarcoma and oesophageal cancer is the combined population figure for males and females while prostate cancer denominator is male population only or half the population of Kaposi’s sarcoma. The smaller the denominator, the higher the mortality rate if the numerators remain equal.

### Association between cancers and HIV in Zambia

Most cancers in Zambia are associated with HIV or HIV defining malignancies. HIV positive patients are more susceptible to cancers due to their compromised immunity. Results of the linear regression analysis and Pearson correlation indicated a strong association between cancer and HIV with *p*-value of 0.000 and correlation coefficient of 0.818. Cancer cases, HIV positive cases and percentage of HIV positive cancer cases are shown in **Table 6.**

**Table 6:** Association between cancers and HIV in Zambia, 2007 – 2014

| ICD10 | Cancer Type | Cases | HIV positive | Percentage of HIV + | *r* | *p*- value |
| --- | --- | --- | --- | --- | --- | --- |
| **C00-C06** | Oral Cavity | 169 | 20 | 12 |  |  |
| **C09-C10, C12-C14** | Other Pharynx | 42 | 4 | 10 |  |  |
| **C11** | Nasopharynx | 85 | 22 | 26 |  |  |
| **C15** | Oesophagus | 581 | 71 | 12 |  |  |
| **C16** | Stomach | 395 | 41 | 10 |  |  |
| **C18-C21** | Large Bowel | 539 | 60 | 11 |  |  |
| **C22** | Liver | 460 | 43 | 9 |  |  |
| **C25** | Pancreas | 102 | 6 | 6 |  |  |
| **C32** | Larynx | 125 | 14 | 11 |  |  |
| **C33-C34** | Lung | 135 | 12 | 9 |  |  |
| **C40-C41** | Bone | 252 | 6 | 2 |  |  |
| **C43** | Melanoma of the skin | 145 | 11 | 8 |  |  |
| **C44** | Other Skin | 406 | 65 | 16 |  |  |
| **C46** | Kaposi's Sarcoma | 2891 | 1734 | 60 |  |  |
| **C50** | Breast | 1469 | 169 | 12 |  |  |
| **C51-C52** | Vulva/Vagina | 246 | 58 | 24 |  |  |
| **C53** | Cervix Uteri | 7389 | 1331 | 18 | **0.818** | **0.000** |
| **C54** | Corpus Uteri | 95 | 8 | 8 |  |  |
| **C55** | Uterus | 115 | 14 | 12 |  |  |
| **C56** | Ovary | 238 | 25 | 11 |  |  |
| **C60** | Penis | 166 | 29 | 17 |  |  |
| **C61** | Prostate | 1651 | 99 | 6 |  |  |
| **C64** | Kidney | 171 | 4 | 2 |  |  |
| **C67** | Bladder | 478 | 22 | 5 |  |  |
| **C69** | Eye | 909 | 154 | 17 |  |  |
| **C70-C72** | Brain, Nervous System | 126 | 10 | 8 |  |  |
| **C73** | Thyroid | 92 | 8 | 9 |  |  |
| **C81** | Hodgkin Disease | 151 | 31 | 21 |  |  |
| **C82,C85,C96** | Non-Hodgkin Lymphoma | 665 | 161 | 24 |  |  |
| **C90** | Myeloma | 45 | 3 | 7 |  |  |
| **C91-C95** | Leukaemia | 126 | 5 | 4 |  |  |
| **C07-C08, C17,...** | Others | 1054 | 97 | 9 |  |  |
|  | **Totals** | **21512** | **4337** | **20** |  |  |

## DISCUSSION

Although cancers are widely distributed in Zambia, this study has established that cancers are more prevalent in the southern zone than the northern zone. The southern zone comprises the high prevalence region and medium prevalence region. Lusaka and Eastern provinces with very high cancer prevalent districts such as Luangwa, Katete, Chipata and Lusaka form the belt of the high prevalence region while Central, Southern and Western provinces with high cancer prevalent districts such as Kabwe, Mongu and Monze among others form the medium prevalence region. Although the northern zone has relatively low prevalence of cancers, there are notable districts such as Kasempa, Kasama, Ndola and Mansa with relatively high prevalence rates.

Some studies argue that the observed geographic variation in cancer distribution is due to differences in the availability of cancer screening and detection facilities [24-26]. Our study however has established that geographiclocation plays a big role in the pattern of cancer distribution in Zambia. This observation was made after mapping cancer cases based on patients’ residential addresses. Distinct patterns were geographically displayed implying that geographic distribution plays a role. Cultural practices and lifestyles are other contributing factors to the observed trends of cancer distribution in Zambia. For instance, cervical cancer which is the most prevalent cancer accounting for 34.3% of all cancers in Zambia has been perpetrated by the culture of polygamist marriages (multiple sexual partners) which has been more pronounced in the southern zone than the northern zone. This culture has led to the spread of human papilloma virus (HPV) which causes cervical cancer [24].

Our survey conducted between October 2017 and February 2018 across Zambia clearly showed that the southern zone of Zambia practices polygamist marriages (multiple sexual partners) far more than the northern zone hence the high prevalence of cervical cancer in the southern zone.

Similar trends in younger generations of European women have been observed and are thought to be due to increasing prevalence of high-risk HPV infection due to changing sexual practices [27,28].

By sex, cancers in Zambia affect more females than males in the ratio 64.86% to 35.14% (F/M = 1.85). This is mostly because of the high prevalence of cervical cancer in Zambian women. This study identified cervical cancer, prostate cancer, breast cancer and Kaposi’s sarcoma as the top four most prevalent cancers in Zambia. Cervical cancer, with standardised incidence rate of 186.3 per 100,000 females has been the leading cause of morbidity and mortality among cancers in Zambian women. This is contrary to other studies which established that cervical cancer is the third leading cause of cancer-related deaths in developing countries which is not the case with Zambia [29]. According to age specific rates of cancers (see Table 4), cervical cancer peak has been observed in the age range of 40 – 49 years implying that this age group is the high risk group for cervical cancer in Zambia.

The prevalence of HPV infection has been 5% in North America and 21% in Africa [24] implying that the risk of contracting HPV in Africa is 4 times higher than in North America. In sub-Saharan Africa, cervical cancer incidence hasalso beeninfluenced by the high prevalence of HIV infection, which has been found to promote progression of cancerous lesions [30]. Cervical cancer incidence rates have decreased by as much as 4% annually, and 70% overall in developed countries where screening programs were introduced several decades ago [31,32]. The observed high prevalence rates of cervical cancer in Zambia is similar to observed trends in Zimbabwe, Uganda and some countries of Central and Eastern Europe [27,33,34].

Prostate cancer has beenthe second most prevalent cancer after cervical cancer and number one cancer in Zambian men with standardised incidence rate of 60.03 per 100,000 males. These results are similar to results of other studies which indicated that prostate cancer is the most prevalent cancer in men [29]. In this study, prostate cancer peak has been observed in the age range of 60 – 69 years. These results indicate that prostate cancer mostly occurs in ageing men. Age is a risk factor for prostate cancer.

Breast cancer ranks third among cancers in Zambia and second in women after cervical cancer. It has a standardised incidence rate of 38.08 per 100,000 females. Breast cancer peak has been observed in the age range of 40 – 49 years which is the productive age group for women; a double impact with cervical cancer in Zambia. On the global scale, breast cancer is the leading cause of cancer-related deaths among females. It is highest in the United States and Western Europe while Africa and Asia have low rates [29]. This explains why breast cancer is ranked third in Zambia. Higher breast cancer incidence in developed countries reflects the use of breast cancer screening as well as higher prevalence of breast cancer risk factors [35]. Risk factors for breast cancer include weight gain after age 18 years, excess body weight (for postmenopausal breast cancer), use of menopausal hormone therapy (MHT), physical inactivity, alcohol consumption, and reproductive and hormonal factors, such as a long menstrual history, recent use of oral contraceptives, and nulliparity or later age at first birth [36,37]. On the other hand, breastfeeding decreases the risk of breast cancer [37].

The distribution of some cancers in Zambia follows the trend of HIV epidemiology. This study has established a strong positive association between cancers and HIV with *p*-value of 0.000 and Pearson correlation coefficient of 0.818. These results indicate that HIV is a risk factor of cancers in Zambia. Some cancers such as Kaposi’s sarcoma are HIV defining malignancies. Kaposi’s sarcoma ranks fourth among cancers in Zambia with standardised incidence rate of 26.18 per 100,000 population. Kaposi’s sarcoma peak has been observed in the age range of 30 – 39 years. This study has observed that Kaposi’s sarcoma is more prevalent in males than in females (see Table 3). The incidence of Kaposi’s sarcoma is higher in Sub-Saharan Africa than in developed countries. The endemic African form of Kaposi’s sarcoma was reported in the 1960s [38], but with the emergence of HIV/AIDS, an atypical aggressive type has been reported in most African countries [39,40]. The incidence of Kaposi’s sarcoma has increased simultaneously with the increased prevalence of HIV[41]. The high prevalence of HIV entails high resultant prevalence of cancers more especially Kaposi’s sarcoma.

Although mortality rate has been high in cervical, prostate and breast cancers as well as oesophageal cancer and Kaposi’s sarcoma, case fatality rate has been moderate in these cancers. This study observed high case fatality rate in Leukaemia, pancreatic cancer, lung cancer and cancers of the brain and nervous system. This means that Leukaemia and pancreatic cancer are more deadly than other cancers, the only holding factor being that their incidence rate is relatively low in Zambia.

This study had some limitations, firstly the missing information in the database made us leave some patients files as they could not be linked to districts of origin. Secondly, the Zambia National Cancer Registry has huge backlog of data due to understaffing. This brought about the adjustment of our study from 2007 - 2017 to 2007 to 2014 period. Lastly, registry based studies have challenges to provide precise data as registries data capturing system do not always suit all study designs. ZNCR does not fully capture risk factors for all cancers hence we could not critically determine all risk factors in this study.

In conclusion, the widespread distribution of cancers with high prevalence in the southern zone has been perpetrated by lifestyle and sexual culture as well as geography. The top seven most cancer prevalent districts in Zambia are Luangwa, Kabwe, Lusaka, Monze, Mongu, Katete and Chipata. Cervical cancer, prostate cancer, breast cancer and Kaposi’s sarcoma are the top four most prevalent cancers as well as major causes of cancer related deaths in Zambia although Leukaemia and pancreatic cancer have the highest case fatality rate in Zambia. Most cancers in Zambia are HIV defining malignancies and their upward trend is due to the increase in HIV cases. Changing the lifestyle and sexual culture would greatly help in the prevention of rampant spread of HPV, HIV, HBV and EBVamong others which would result in the reduction of cancer cases in Zambia. Furthermore, intensifying cancer screening and early detection countrywide would help to drastically reduce cancer prevalence in Zambia.

## Authors’ Contributions

MK designed the study, developed and programmed the model and drafted the manuscript.LL approved the model and provided technical support in analyses. MK and HS conceived the study. All authors read and approved the manuscript.

## Acknowledgements

Compliments go tothe Research Ethics Committee of Institute of Urban Environment, Chinese Academy of Sciences, the University of Zambia Biomedical Research Ethics Committee andthe National Health Research Authority of the Republic of Zambia for allowing the researchers to conduct a study in Zambia. The principal researcher is a PhD candidate at the Institute of Urban Environment, university of Chinese Academy of Sciences, 1799 Jimei Road, Xiamen, 361021, PR China under the scholarship of UCAS.

---

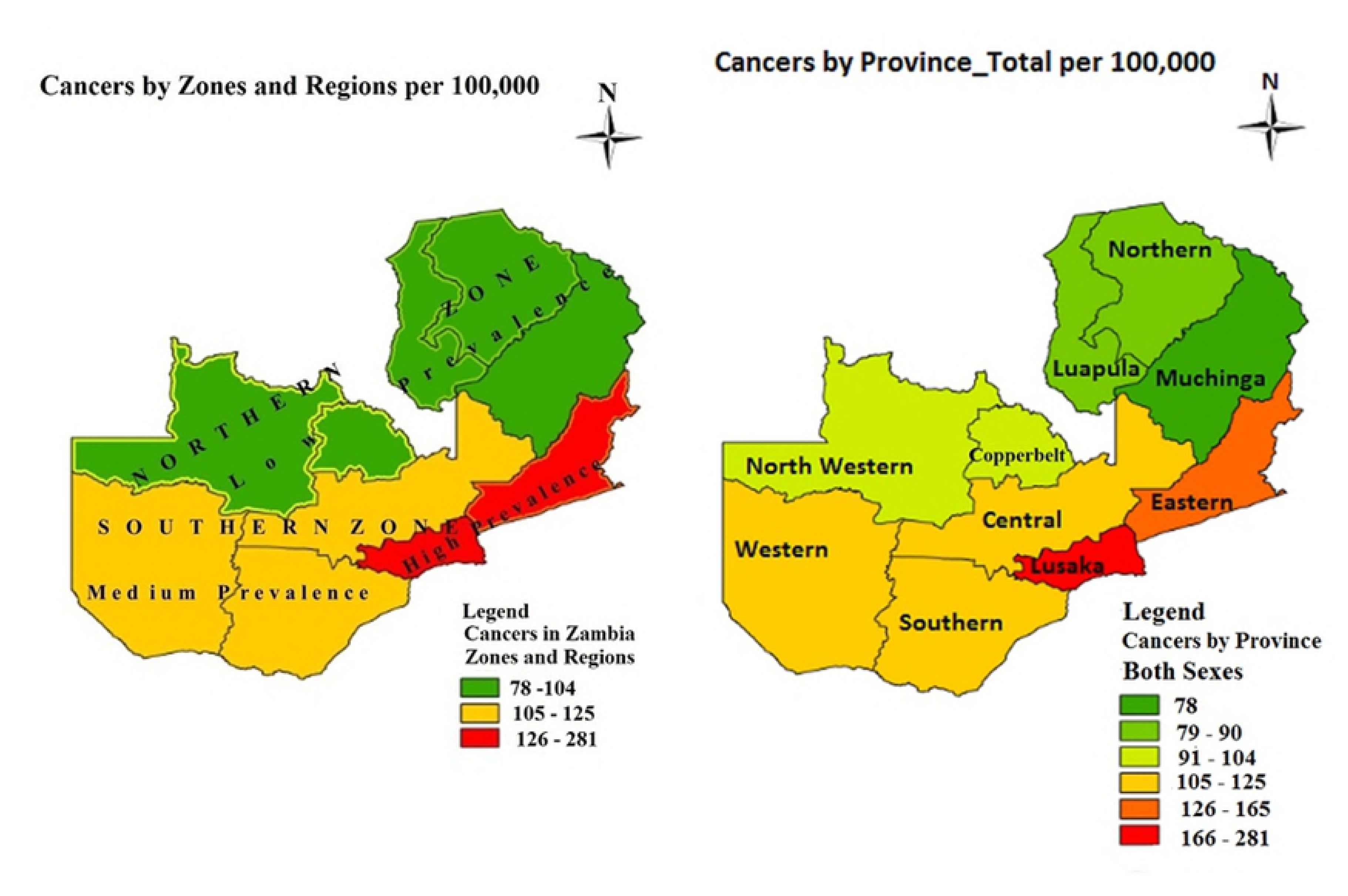

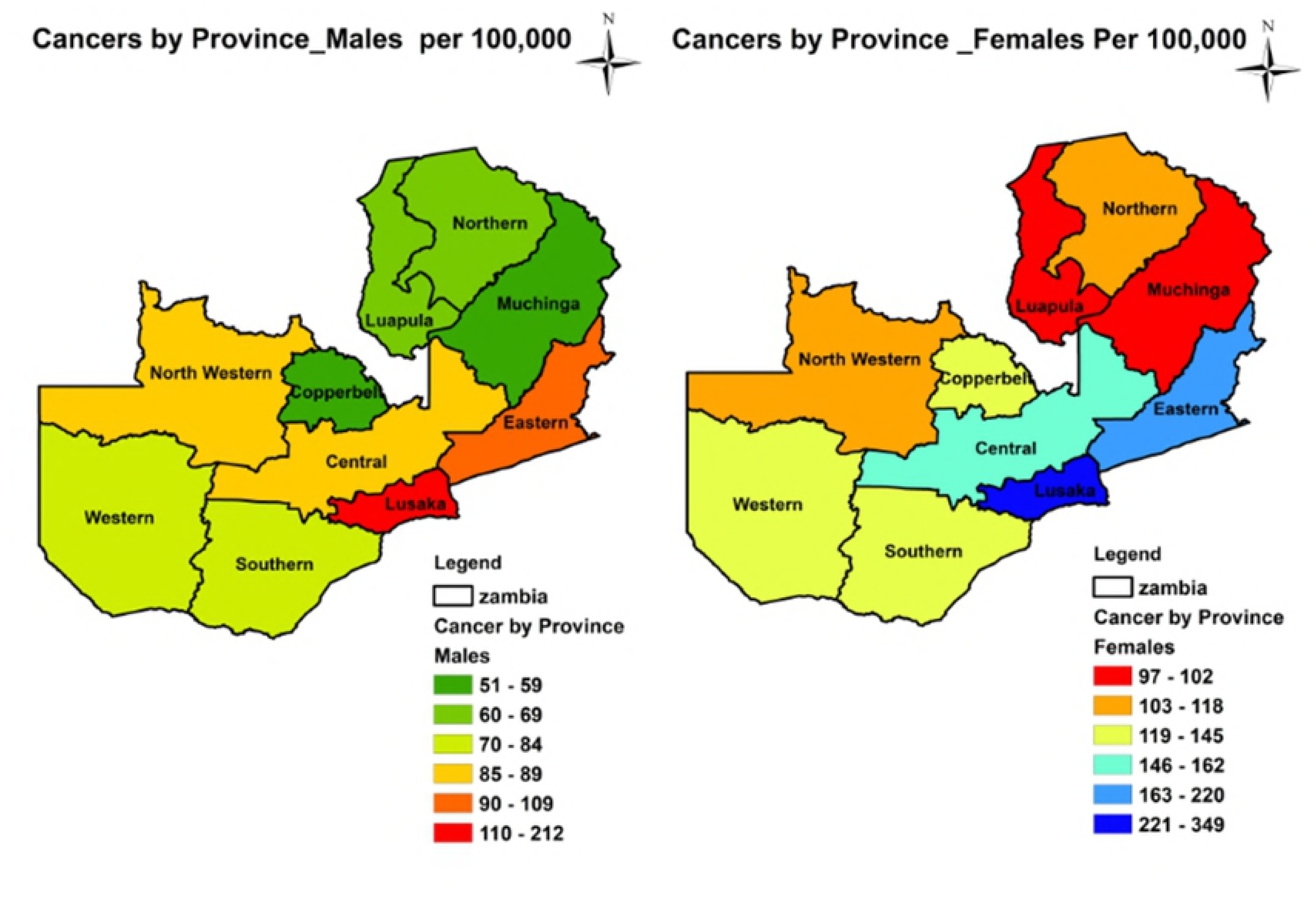

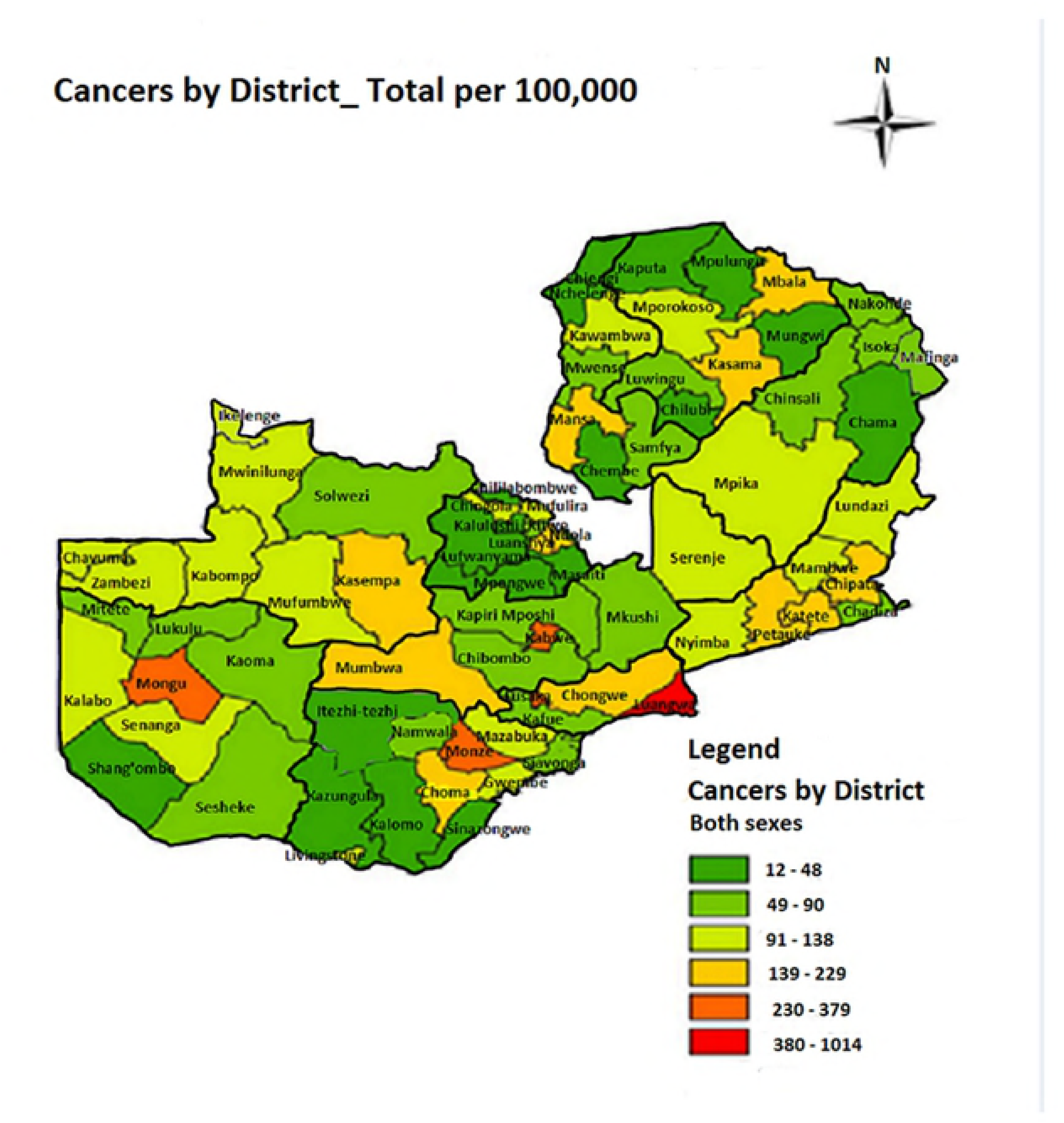

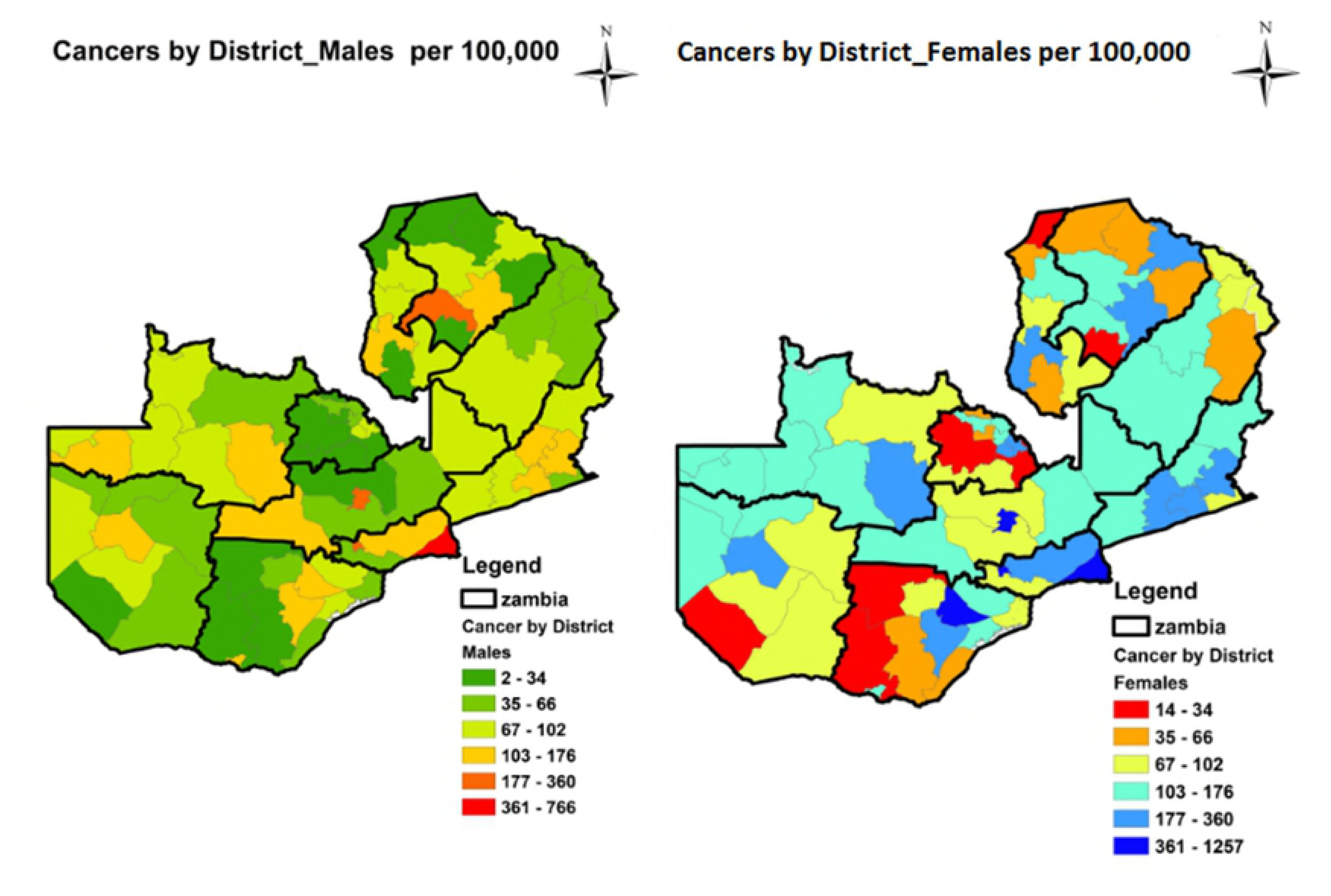

